# Quantitative proteomics and phosphoproteomics reveal glucocorticoid stimulation of TLR and Rho GTPase signaling in neutrophil-like cells

**DOI:** 10.1101/2025.06.25.661639

**Authors:** Hayoung Cho, Michael L. Nielsen, Jesper V. Olsen

## Abstract

**Background:** Glucocorticoids (GCs) are corticosteroid hormones that are commonly used for treating systemic inflammatory diseases and acute infections. Immunosuppressive effects of GCs have been studied in many cell types, particularly macrophages and T cells. Despite the importance and abundance of neutrophils in the human immune system, GC responses remain understudied in neutrophils.

**Results:** Here, we performed quantitative mass-spectrometry (MS)-based proteomics of neutrophil-like cells (NLCs) differentiated from human HL-60 promyelocyte cells. Proteome and flow cytometry analysis showed that NLCs share features of neutrophils. Quantitative proteomics and phosphoproteomics of NLCs treated with two synthetic GC compounds, the clinical drugs dexamethasone (DEX) and prednisolone (PRED), identified higher numbers of significantly regulated proteins and phosphosites compared to parental HL-60 cells. GC treatments triggered abnormal neutrophil activation and aging by promoting toll-like receptor (TLR) signaling and CXCR4 serine phosphorylation. We also identified RIPOR2 as a novel target protein of GC which stimulates Rho GTPase signaling networks and upregulates actin cytoskeletal proteins.

**Conclusions:** Our results not only reveal unconventional regulatory mechanism of GCs in the human immune system but also provide valuable resources for discovering novel GC-responsive protein targets.

## Background

The human immune system plays a vital role in maintaining homeostasis by responding to internal and external stimulants through complex defensive mechanisms against infections and pathogens (1). Advancements in biomedical approaches such as monoclonal antibodies, imaging techniques, genetically engineered animal models, and omics methods have improved our understanding of innate and adaptive immune responses, facilitating new strategies to overcome inflammatory and autoimmune disorders (2).

Neutrophils, also known as the “Cinderella” of the innate immune system (3), are the most abundant cell type among leukocytes and act as the first-line defense against infections to stimulate adaptive immune responses. Neutrophil dysfunctions can lead to serious disorders such as inflammatory bowel disease, rheumatoid arthritis, and multiple sclerosis. Neutrophils are short-lived immune cells with the longest reported *in vivo* lifespan of up to 5.4 days (4), however, this remains debated as older *in vivo* studies suggested significantly shorted lifespans of approximately 10-18 hours (5), and 7-9 hours *ex vivo* (6). Neutrophils are generated and matured in bone marrow and subsequently released to blood circulation, where they sense inflammatory signals through cell surface receptors such as toll-like receptor 2 (TLR2), TLR4, and formyl-peptide receptors (FPRs). Upon receiving inflammatory signals, cells undergo neutrophil extravasation, also known as neutrophil trans-endothelial migration to exit the bloodstream, and migrate to the site of infection through a multistep process of rolling, adhesion, crawling, and transmigration (7). At inflammation sites, activated neutrophils migrate towards infected tissues through neutrophil chemotaxis where they combat pathogens by deploying degranulation, phagocytosis, and release of neutrophil extracellular traps (NETs), which are mesh-like structures composed of DNA, histones and granular proteins to trap and kill pathogens. Remaining mature neutrophils in the bloodstream undergo aging that is primarily driven by TLR-Myd88 signaling and internal circadian clock cues (8). Aged neutrophils are cleared from circulation to bone marrow and other tissues for balanced level of circulating neutrophils (9, 10).

Extravasation, chemotaxis, and NET release are unique features of neutrophils that accompany dynamic cytoskeletal remodeling maintained by crosstalk among Rho GTPases. Rho GTPases, such as Rac, Rho, and Cdc42, are small proteins of the Ras superfamily that are activated upon GTP binding. Rho GTPases are key signaling molecules for adaptive cytoskeletal rearrangements, cell motility, and cell polarity (11). In neutrophils, Rac promotes protrusion of the leading edge through actin polymerization while RhoA promotes myosin contraction at the trailing edge. Cdc42 regulates microtubule stabilization and F-actin polymerization, supporting neutrophil migration towards chemokines through diverse cytoskeletal remodeling at both edges (12, 13). As a result, Rho inhibition suppresses AKT activation, reducing neutrophil infiltration and NET release in mice exposed to immunostimulants (14).

Glucocorticoids (GCs) are a type of corticosteroid hormone that primarily binds to glucocorticoid receptor (GR), encoded by the *NR3C1* gene. GR is a prominent member of the nuclear receptor superfamily that translocates from cytoplasm to nucleus upon GC binding and functions as a transcription factor regulating inflammatory signaling genes (15). Due to their significant effects on pro- and anti-inflammatory genes through direct transcriptional and non-genomic regulations, GCs are commonly used to treat chronic and acute inflammations (16). In addition, solid malignant cancer patients are often treated with synthetic GCs to alleviate the side effects of chemotherapy (17). Recently, systemic administration of GCs effectively treated severe COVID-19 (18). To understand the multifaceted functions and potential side effects of GC treatments, omics approaches have been extensively used to unveil GC actions (19). However, further investigations of GC mechanisms in neutrophils are necessary to uncover the signaling pathways driving their responses that are contradictory from well-established immunosuppressive mechanisms. For example, the cellular signaling underlying the increase of circulating neutrophil counts in GC-treated patients, a phenomenon known as GC-induced leukocytosis, are not yet fully understood (20). Moreover, a recent study suggests the potential adverse effects of GCs by demonstrating that GCs create metastasis-friendly tumor microenvironment *in vivo* by promoting neutrophils to release NETs (21). Neutrophils are terminally differentiated cells with limited transcriptional activity (9). Therefore, quantitative proteomics strategy can be highly beneficial to study the molecular complexity and dynamics of neutrophils through protein abundance and post-translational modifications (PTMs) (22).

In this study, we used cutting-edge mass spectrometry (MS) techniques to study the dynamic signaling responses of human neutrophils to GCs by acquiring deep proteome and phosphoproteome of neutrophils differentiated from human promyelocyte HL-60 cell line. To our understanding, this is the first proteomics study on GC-treated human neutrophils. Our large-scale quantitative dataset contributes to better understanding of GC systemic regulatory responses through identification of GC-sensitive biomarkers and post-translational signaling pathways.

## Results

### Fig. 1: DMSO differentiated HL-60 cells resemble neutrophil-like characteristics

To confirm the effects of non-specific activation of neutrophils in 2D mono-culture (23), we compared the proteomes of neutrophils lysed immediately after isolation from human buffy coats with isolated neutrophils lysed after 2 hour incubation in 0.1% dimethyl sulfoxide (DMSO), a common solvent used with cells for cryopreservation and as a solvent for drugs (24)(Fig.1A). Following DMSO incubation, proteins associated with Reactome pathway of extracellular matrix organization (HSA-1474244) were significantly upregulated and proteins enriched in eukaryotic translation termination (HSA-72764) were significantly downregulated by a fold-change of at least two (Fig.1B). To overcome the limitation of the short lifespan and atypical activation of neutrophils, we performed quantitative mass-spectrometry (MS) analysis on neutrophil-like cells (NLCs) differentiated from HL-60 cells by incubation in 1.25% DMSO for four days (Fig.1A). Comparing the proteomes of HL-60 cells, NLCs, and neutrophils showed an overlap of 5981 protein groups, which accounted for 71%, 72%, and 92% of the total proteome, respectively (Fig.1C). The 144 protein groups only quantified in NLCs and neutrophils included neutrophil activation surface markers (CD62L, CD66a, and CD66b) (25, 26), and pathogen recognition proteins (FPR1, FPR2, TLR4, and S100A12) (27, 28). These unique proteins showed Reactome pathway enrichments in neutrophil degranulation (HSA-6798695) and innate immune system (HSA-168249) (Fig.S1A). This analysis suggested that the differentiated NLCs are characterized by specifically expressing the important neutrophil-specific cell surface receptors and signaling pathway components.

**Figure 1.**
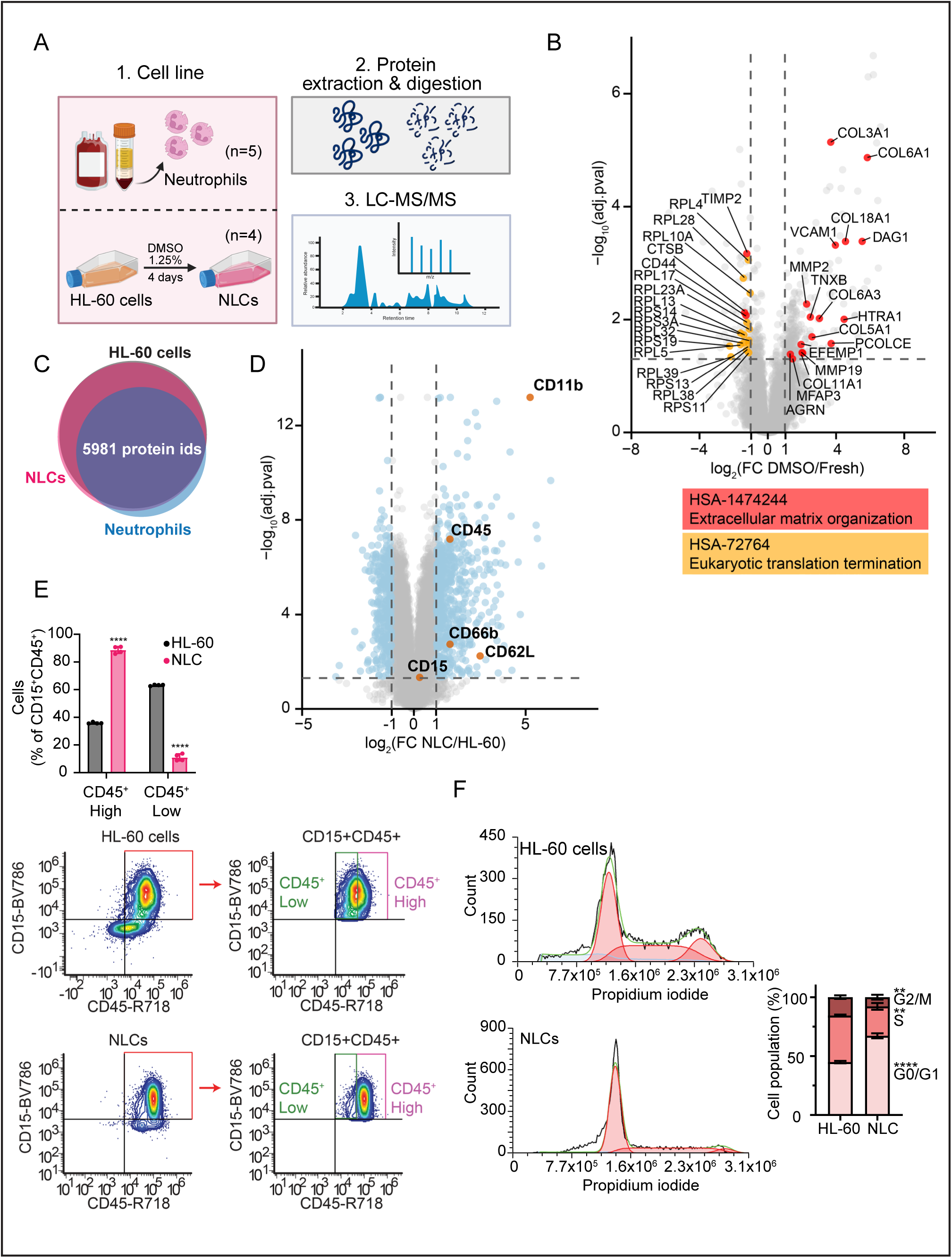
Neutrophil phenotype characterization in NLCs. A) Schematic illustration of proteomic analysis in neutrophils isolated from human blood and NLCs differentiated from HL-60 cells. Illustration was created with Biorender.com. B) Volcano plot comparing protein expressions of neutrophils lysed after 2 hour DMSO incubation versus freshly lysed after isolation. Differential expression analysis was performed using one-way ANOVA with Tukey’s HSD correction. Red and yellow dots represent proteins with statistically significant differences (adjusted *p*-value < 0.05 and log_2_FC > |1|), enriched in Reactome pathway of extracellular matrix organization and eukaryotic translation termination, respectively. C) Venn diagram showing the overlaps among quantified proteins in proteome of HL-60 cells, NLCs, and neutrophils. D) Volcano plot comparing protein expressions of NLCs versus HL-60 cells using one-way ANOVA with Tukey’s HSD correction. Blue dots represent proteins with statistically significant differences (adjusted *p*-value < 0.05 and log_2_FC > |1|) and orange dots represent mature neutrophil markers. E) CD15 and CD45 staining analysis of HL-60 cells and NLCs using flow cytometry. CD45^+^Low/High populations are highlighted under CD15^+^CD45^+^ gate. F) Cell cycle arrest assay of HL-60 cells and NLCs with propidium iodide staining using flow cytometry. Histograms display DNA content with bar plots representing G0/G1, S, and G2/M phases. Multiple unpaired t-test with Welch correction was used for flow cytometry analysis.

After differentiation from HL-60 cells, NLCs significantly overexpressed a pan-leukocyte marker (CD45) and neutrophil activation markers (CD11b, CD66b, and CD62L) (29) at least by two-folds. CD15, another neutrophil maturation marker (30), increased by ∼19.5% (Fig.1D). Flow cytometry analysis showed that the total population of CD15^+^CD45^+^ cells remained unchanged after differentiation (Fig.S1B), however, CD15^+^CD45^+^ cells expressed substantially high CD45 level in NLCs compared to HL-60 cells (Fig.1E). Lastly, NLCs were arrested at the G0/G1 phase of the cell cycle (Fig.1F), demonstrating a loss of cell division capacity for neutrophils (31). Together, we provide evidence that human buffy coat-isolated neutrophils are not ideal for *ex vivo* studies and that NLCs differentiated from HL-60 cells exhibit neutrophil-like phenotypes.

### Fig. 2: GC treatments promote significant protein regulations in NLCs through phosphorylation of GR

RNA-sequencing data from the Human Protein Atlas (HPA) suggests that GR expression is the second highest in neutrophils among immune cells and the highest in bone marrow among tissues, highlighting its prominent role in innate immune function (Fig.2A, S2A). To understand the effects of GR regulation on neutrophils, we first measured the cytotoxicity of GC treatment in NLCs. 24 hours of dexamethasone (DEX) treatment dose-dependently suppressed the cell viability of HL-60 cells (Fig.2B). In contrast, neither cell viability nor apoptosis were affected by GC treatments in NLCs (Fig.2C, S2B).

**Figure 2.**
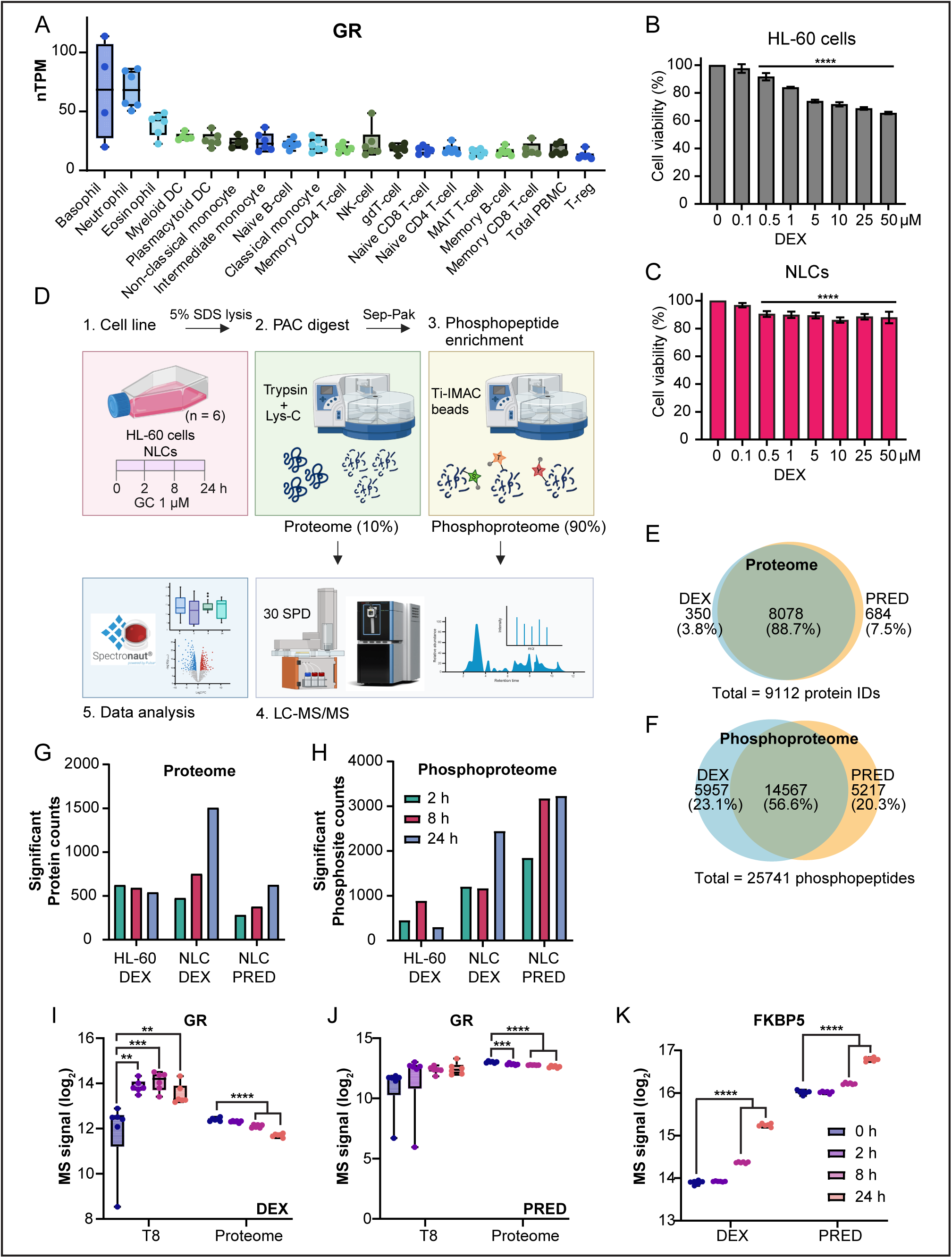
MS-based proteomic and phosphoproteomic analysis of HL-60 cells and NLCs after time-course treatment of GCs. A) GR mRNA levels represented as normalized transcript per million (nTPM) in human immune cell lines. RNA-seq data was acquired from the Human Protein Atlas (HPA). B-C) Cell viability assay of HL-60 cells (B) and NLCs (C) after dose-dependent titration of DEX for 24 hours. D) Schematic illustration of proteome and phosphoproteome workflow after time-course treatment of 1 µM GC treatments in HL-60 cells and NLCs. Spectronaut logo was obtained from Biognosys AG and Orbitrap Astral MS image was obtained from Thermo Fisher Scientific. Illustration was created with Biorender.com. (E-F) Venn diagrams showing overlaps between DEX and PRED treatments in proteome (E) and phosphoproteome (F) profiles. (G-H) Bar plots showing the number of proteins (G) and phosphosites (H) with statistically significant regulations compared to 0 hour control using one-way ANOVA and Tukey’s HSD correction (adjusted *p*-value < 0.05). (I-J) Box plots showing individual log_2_ transformed label-free quantification of GR phosphorylation at Thr-8 and total form after DEX (I) and PRED (J) treatments. (K) Box plots showing log_2_ transformed label-free quantification of FKBP5 protein expression. One-way ANOVA with Dunnette’s multiple comparisons test was used for statistical tests.

To further investigate glucocorticoid responses in an unbiased manner, we performed quantitative MS using two types of corticosteroids, thereby minimizing the risk of drug-specific effects. DEX and prednisolone (PRED) are both FDA-approved corticosteroids used for treating various inflammatory conditions such as allergic reactions, dermatologic conditions, endocrine disorders, and respiratory diseases (32). We acquired deep proteome and phosphoproteome of NLCs and HL-60 cells treated with DEX (1 µM) and PRED (1 µM) to explore the altered GC sensitivity following differentiation (Fig.2D). Across a time-course treatment of 0, 2, 8, 24 hours, we quantified 9112 proteins and 25741 phosphopeptides using a label-free approach. We observed substantial overlap between DEX and PRED treatments, with 88.7% of shared protein IDs and 56.6% of shared phosphopeptides (Fig.2E,F). Next, we evaluated the effects of GCs by protein and phosphosite abundance changes that were statistically significant with adjusted *p*-value of less than 0.05. Interestingly, the number of significantly regulated proteins remained relatively constant over time in the HL-60 proteome, while the number for NLC proteome exhibited a time-dependent increase (Fig.2G). Additionally, the number of significantly regulated phosphosites was considerably higher in NLCs compared to HL-60 cells (Fig.2H). This suggested that the regulatory effects of GCs were more pronounced in NLCs than in HL-60 cells. Following GC treatments, time-dependent increase of GR phosphorylation at Thr-8 and ubiquitin-mediated protein degradation of the total form were detected (33)(Fig.2I,J, Fig.S2C,D). Protein expression of FKBP5, a co-chaperone of HSP90 essential for GR activities (34), was also increased significantly by DEX and PRED treatments (Fig.2K, Fig.S2E). GC treatments increased GR phosphorylation at Ser-211 and decreased the total protein in time- and dose-dependent manners as demonstrated by the immunoblotting analysis (Fig.S2F,G). Collectively, we confirmed that the protein regulations were driven by GC-mediated GR modulation.

### Fig. 3: GC treatments activate NLCs through TLR cascades at early responses

We next investigated whether specific signaling pathways were associated with GC-responsive mechanisms in NLCs. We performed fuzzy c-means clustering using the Mfuzz algorithm (35) to group similar temporal expression profiles of proteins with significant regulation (adjusted *p*-value less than 0.05) into eight distinct clusters. After DEX treatment, clusters 1 and 6 showed increasing expression patterns, with cluster 1 proteins exhibiting early responses from 2 hours and cluster 6 responding later at 24 hours (Fig.3A). Similarly, after PRED treatment, cluster 1 showed early activation at 2 hours and cluster 4 responded from 24 hours (Fig.S3A). Proteins in cluster 1 were significantly enriched in Reactome pathway of TLR cascades (HSA-168898) (Fig.3B, S3B). TLRs are known to activate innate immune responses by recognizing diverse pathogens (36). TLR2 was robustly increased by GC treatments, together with PER1 which is a circadian regulator important for neutrophil release (37) and FPR1, a cell surface receptor essential for neutrophil activation (38) (Fig.3C-F, S3C,D). TLR2 and PER1 increments by DEX treatment were restored to the control level when DEX and mifepristone (MIF), a GR antagonist, were treated in combination, indicating that they are specifically regulated by GCs (Fig.S3E). These results suggest that GC treatments upregulate proteins associated with neutrophil activation.

**Figure 3.**
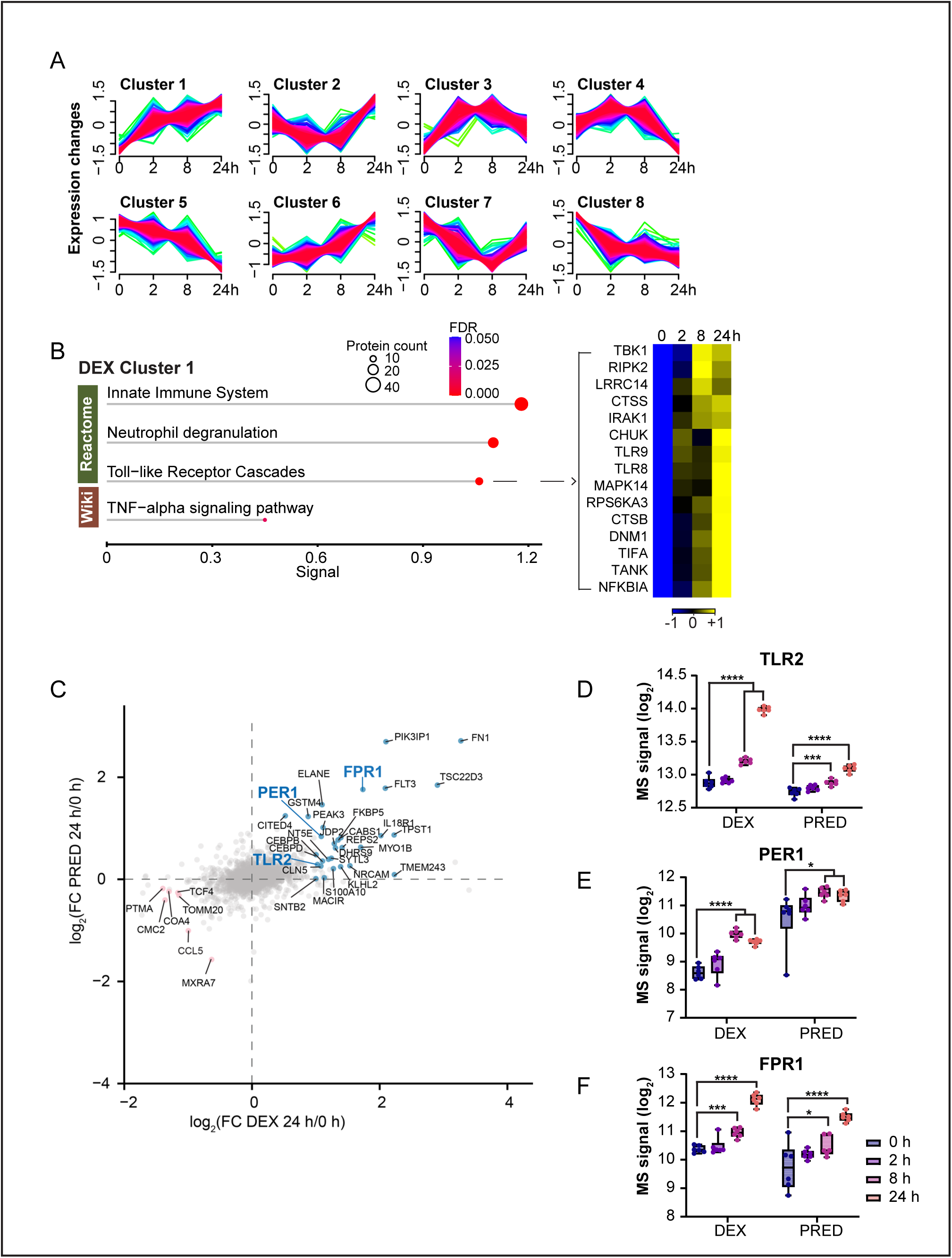
Proteomic analysis of NLCs after time-course treatment of DEX and PRED showing neutrophil activation signals. A) Soft clustering analysis with statistically significant proteins (adjusted *p*-value < 0.05, one-way ANOVA with Tukey’s HSD correction) after DEX treatment in NLCs. Mfuzz clustering package was used with fuzzification factor *m* = 1.5 and cluster number 8. B) Functional enrichment of proteins in cluster 1 using string analysis after DEX treatment. Dot size represents the number of proteins and dot color represents FDR values. Heatmap represents proteins in TLR cascades with z-scored log_2_ transformed label-free quantification, colors showing z scores between −1 and 1. C) Scatter plot comparing protein expression changes after 24 hours treatment of DEX (x-axis) and PRED (y-axis) relative to 0 h control. Labeled dots represent proteins significantly regulated (adjusted *p*-value < 0.05 and log_2_FC > |1|) after one-way ANOVA with Tukey’s HSD correction in either DEX or PRED treatment. D-F) Box plots showing individual log_2_ transformed label-free quantification of TLR2 (D), PER1 (E), and FPR1 (F) protein expressions after DEX and PRED treatments. One-way ANOVA with Dunnette’s multiple comparisons test was used.

### Fig. 4: GC treatments induce abnormal neutrophil activation and aging by MAPK cascade

Based on previous reports that the mitogen-activated protein kinases (MAPK), ERK and p38MAPK, are key signaling pathways of FPR activation and migration in neutrophils (39, 40), we investigated the effects of GCs on protein phosphorylation dynamics by immunoblotting and quantitative phosphoproteomics. We confirmed that the activation loop phosphorylation sites of ERK1/2 and p38MAPK, but not AKT, showed time- and dose-dependent increases after GC treatments. Furthermore, increased Rac2 and constant expression of Ras suggested that GC treatments activated MAPK-associated pathways downstream of Ras (Fig.4A,B). From the phosphoproteome data, we observed a prominent increase in the activating Thr-735 phosphorylation of ADAM17, which is regulated by ERK1/2 (41) and p38MAPK (42) (Fig.4C, S4A). Upon neutrophil activation, active ADAM17 cleaves CD62L for neutrophil recruitments (43). These data support our finding that GC promotes neutrophil activation and it is driven by dynamic MAPK signaling.

**Figure 4.**
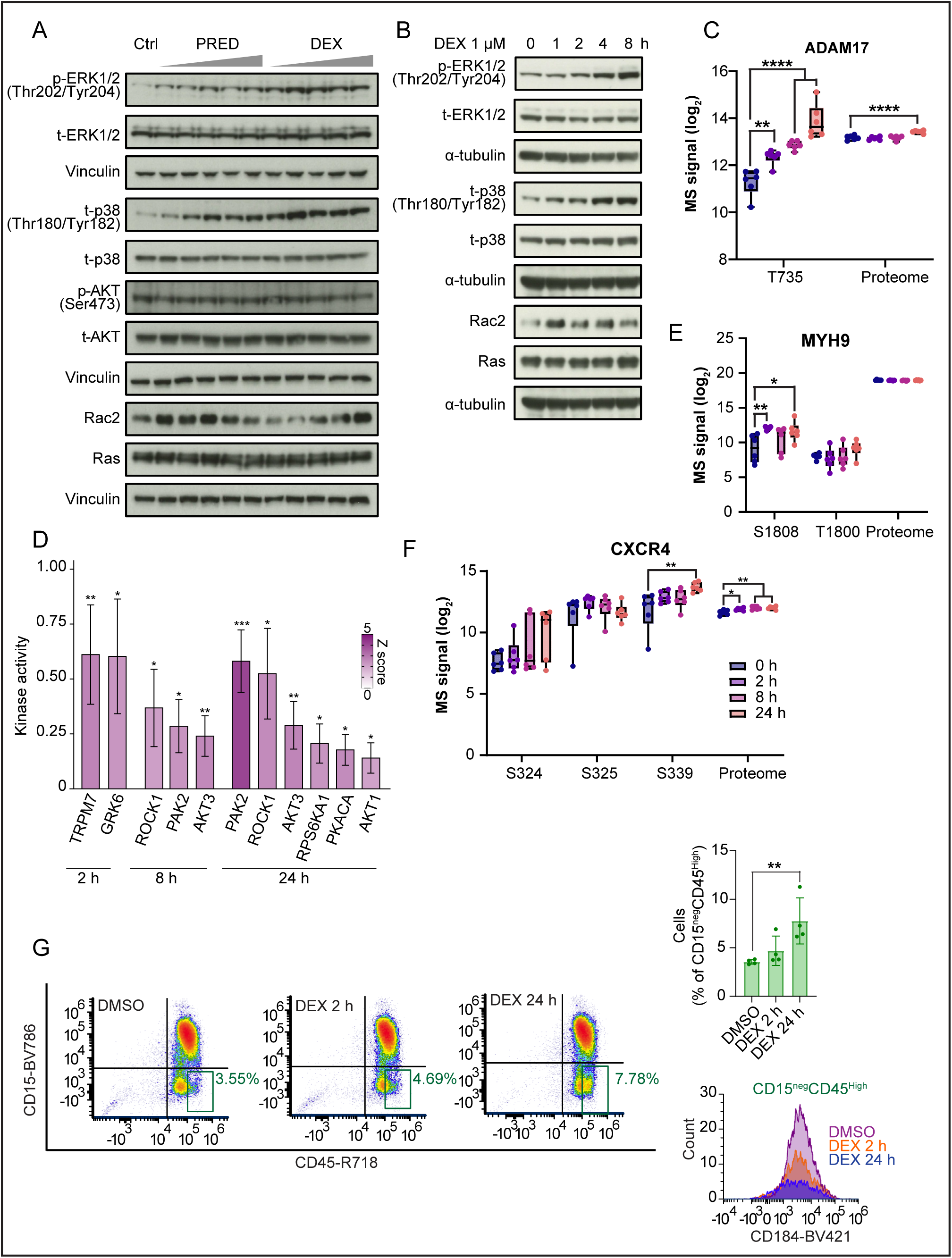
Investigation of DEX treatment effects driving neutrophil activation and aging. A) Immunoblotting analysis of MAPK signaling proteins after 24 hours dose-dependent treatment with PRED and DEX (0.1, 0.5, 1, 5, 10 µM). B) Immunoblotting analysis of MAPK signaling proteins after time-dependent treatment of 1 µM DEX (1, 2, 4, 8 hours). C) Box plots showing individual log_2_ transformed label-free quantification of ADAM17 phosphorylation at Thr-735 and protein expression. D) Kinase activity analysis using RoKAI app after treatment of DEX. Bar color represents z-scored kinase activity (0 to 5). E-F) Box plots showing individual log_2_ transformed label-free quantification of phosphorylation and protein expressions of MYH9 (E) and CXCR4 (F) after DEX treatment. G) Flow cytometry analysis of 2 and 24 hours DEX treatments in NLCs. Cells were stained with CD15, CD45, and CXCR4 (CD182). CD15^neg^CD45^High^ cell populations are highlighted under live cell gate. Bar plot shows the number of CD15^neg^CD45^High^ cells after DEX treatment and histogram represents CXCR4 expression of CD15^neg^CD45^High^ cells. One-way ANOVA with Dunnette’s multiple comparisons test was used.

Next, we employed the RoKAI algorithm (44) for inferring kinase activities from the time-course treated phosphoproteome. After 2 hours of DEX treatment, TRPM7 and GRK6 kinases were activated (Fig.4D). We used kinase-substrate data obtained from PhosphositePlus and SIGNOR 3.0 to identify direct substrates of TRPM7 and GRK6. TRPM7 increased phosphorylation of MYH9 Ser-1808 and Thr-1800 (Fig.4E, S4B), which are associated with cell migration and translocalization of the chemokine receptor CXCR4 to nucleus (45, 46). Following DEX treatment, GRK6 phosphorylation targets included CXCR4 at Ser-324, Ser-325, and Ser-339 (Fig.4F, S4C). CXCR4 phosphorylation was also significantly increased at Ser-324 and Ser-325 after PRED treatment (Fig.S4D,E). Phosphorylation at Ser-324 and Ser-325 is critical regulator of CXCR4 desensitization, β-arrestin recruitment, and internalization of CXCR4 (47). Additionally, aged neutrophils feature significant upregulation of CXCR4 (48) and GRK6 kinase activity regulates the number of circulating neutrophils in mouse (49). We then performed flow cytometry to investigate cell activation and maturity. DEX treatment increased the number of immature NLCs, identified as CD15^neg^CD45^high^ cells, which exhibited high CXCR4 (CD184) (Fig.4G). Collectively, phosphorylation analysis supports our finding that DEX treatment drives abnormal cell activation and aging by promoting MAPK signaling pathways.

### Fig. 5: GCs promote Rho GTPase signaling pathway and actin dynamics by RIPOR2 phosphorylation at later responses

After 24 hours of GC treatments, significantly upregulated proteins in cluster 6 (DEX) and cluster 4 (PRED) were associated with Reactome pathway of Rho GTPase cycle (HSA-9012999, HSA-9013026, and HSA-9013106) (Fig.5A, S5A). This result corresponded to the RoKAI kinase activity analysis that showed ROCK1 and PAK2 activation after 8 and 24 hours following DEX treatment (Fig.4D). This also led to significant upregulation of ARHGAPs, which are direct substrates of ROCKs and PAKs involved with actin dynamics (50–52) (Fig.S5B). We also detected decreased Tyr-291 phosphorylation of WASP, another downstream molecule of Rho GTPases (Fig.S5C,D). This site is known to regulate WASP activity in cytoskeletal rearrangements, indicating GC-regulation through WASP of actin polymerization (53). As a result, actin cytoskeletal proteins, including MYO1B, ELANE, THBS1, FN1, MFGE8, KLHL2, and PEAK3, were significantly increased after GC treatments (Fig.3C, 5B).

**Figure 5.**
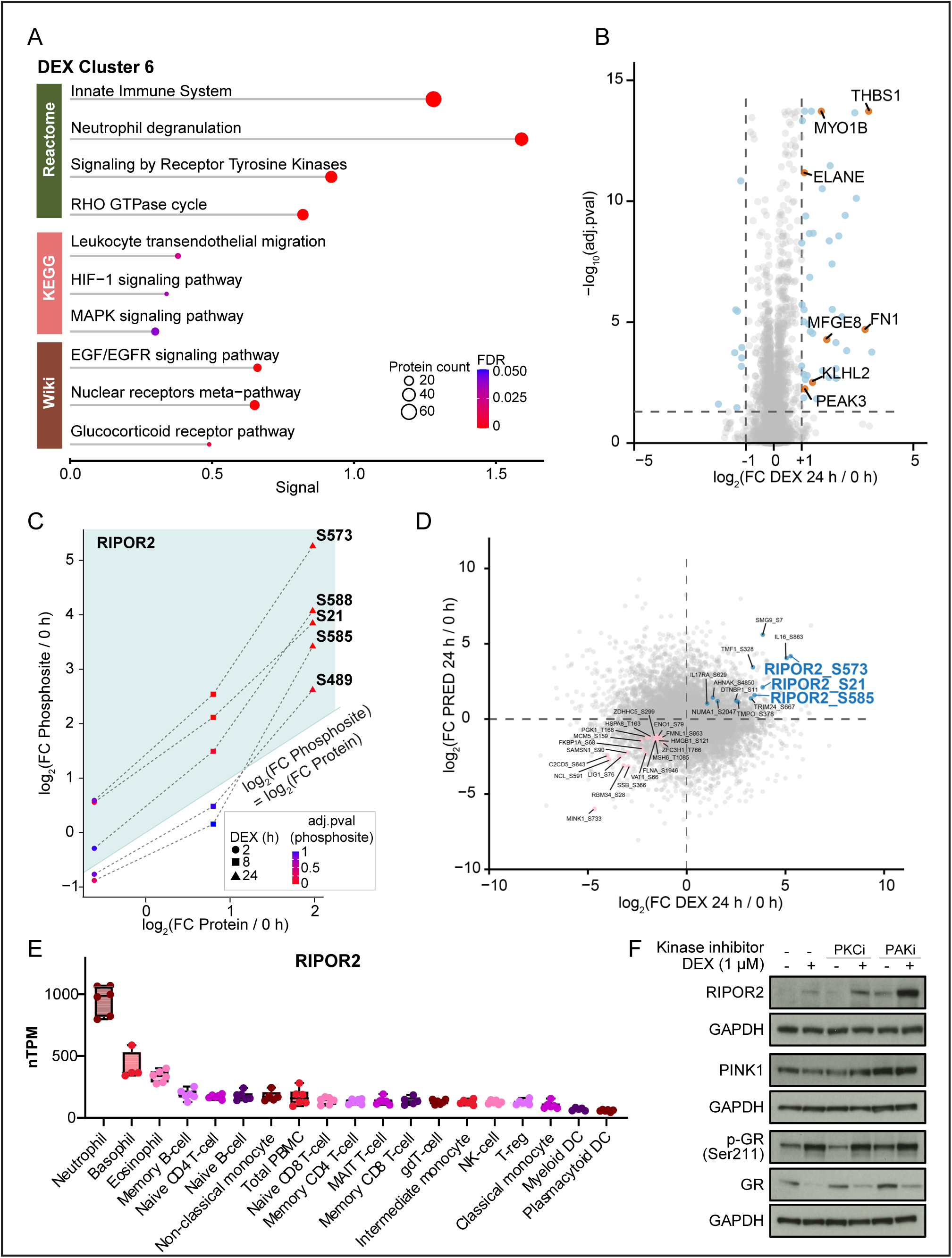
Investigation of DEX treatment effects in Rho GTPase signaling pathway. A) Functional enrichment of proteins in cluster 6 using String analysis after DEX treatment. Dot size represents the number of proteins and dot color represents FDR values. B) Volcano plot comparing protein expressions of DEX 24 hours treatment versus DMSO control in NLCs using one-way ANOVA with Tukey’s HSD correction. Blue dots represent proteins with statistically significant differences (adjusted *p*-value < 0.05 and log_2_FC > |1|) and orange dots represent proteins associated with actin cytoskeleton. C) Line plot comparing RIPOR2 protein expression changes (x-axis) and serine phosphorylation changes (y-axis) after DEX treatment in NLCs compared to 0 hour control. DEX treatment timepoints are represented with different symbols (circle: 2 hours, square: 8 hours, triangle: 24 hours) and symbol colors represent adjusted *p*-value of phosphoproteome after one-way ANOVA with Tukey’s HSD correction. Phosphosites above the blue line show greater increases than the protein expression. D) Scatter plot comparing phosphorylation level changes after 24 hours treatment of DEX (x-axis) and PRED (y-axis) relative to 0 h control. Labeled dots represent phosphosites significantly regulated (adjusted *p*-value < 0.05 and log_2_FC > |1|) after one-way ANOVA with Tukey’s HSD correction in both DEX and PRED treatment. E) RIPOR2 mRNA levels represented as normalized transcript per million (nTPM) in human immune cell lines. RNA-seq data was acquired from the Human Protein Atlas (HPA). F) Immunoblotting analysis of PKCi (Sotrastaurin) and PAKi (PF-3758309) in combination with DEX treatment for 24 hours in NLCs.

Noticeably, we found that RIPOR2 phosphorylation was increased at multiple serine sites (21, 489, 573, 585, 588) after DEX treatment. RIPOR2 is one of the proteins that were specifically regulated by DEX treatment (Fig.S3E), and we confirmed that phosphorylation upregulations were not driven by the increased level of total RIPOR2 protein (Fig.5C, S5E,F). Phosphorylation at Ser-21, Ser-573, and Ser-585 were robustly increased by both DEX and PRED treatments, strongly indicating that RIPOR2 phosphorylation is significantly regulated by GCs (Fig.5D, S5G,H). RIPOR2 mRNA levels were the highest in neutrophil among immune cell lines and in lymph node among tissues from HPA dataset, highlighting a clinical significance of RIPOR2 in innate immunity (Fig.5E, S6A). Furthermore, RIPOR2 is an essential protein for neutrophil directional polarization that exerts RhoA suppressive function by accumulating at the leading edge, and its function is lost by phosphorylation (54).

PINK1 is the only known kinase for RIPOR2 (55) and its transcription is reduced by the phosphorylation of FOXO3a Thr-32, which is an inactive form driven by PI3K/AKT pathway (56). Although PINK1 was not quantified in our MS datasets, activation of AKT from RoKAI analysis and a significant increase in FOXO3 Thr-32 phosphorylation following DEX treatment provide evidence that RIPOR2 phosphorylation is possibly independent of PINK1 (Fig.4D, S6B,C). To better understand the upstream kinases regulating RIPOR2, we quantified RIPOR2 protein levels after treating NLCs with several kinase inhibitors in combination with DEX. AKTi (AKT-1/2i and MK-2206), PKCi (Bisindolylmaleimide I and Sotrastaurin) and PAKi (PF-3758309) were selected based on previous studies that AKT, PKC, and PAK are potential kinases of RIPOR2 (57). Additionally, we tested the effects of FLT3i (KW-2449 and Gilteritinib) based on the robust increase in FLT3 levels after GC treatment in our proteome data (Fig.3C, S3E). AKTi and FLT3i had no effect on RIPOR2 expressions, whereas substantial increases were observed after PKCi (Sotrastaurin) and PAKi (PF-3758309) treatments. Interestingly, single treatment of PF-3758309 induced phosphorylation of GR at Ser-211, which potentially explains a synergistic increase of RIPOR2 after the combination treatment with DEX. Furthermore, PF-3758309 treatment increased PINK1 expression, which was decreased by DEX treatment (Fig.5F). Overall, PAK inhibition promoted GR phosphorylation necessary for GC-mediated RIPOR2 regulation, suggesting that DEX regulates Rho GTPase signaling and actin dynamics through a PAK-RIPOR2 signaling axis.

## Discussion

In this study, we describe for the first time the proteome and phosphoproteome analyses of GC regulations in human NLCs. Our data demonstrates that synthetic GC treatments promote dysfunctional activation and aging of NLCs in absence of pathogens. From proteome profiling, we found significant upregulation of the pattern recognition receptors, TLR2 and FPR1, and circadian clock protein PER1. Furthermore, phosphoproteome analysis exhibited a remarkable increase of CXCR4 phosphorylation and activated MAPK-associated kinases. Our data support a recent finding that GCs induce chronic stress *in vivo* and alters diurnal aging process in mouse neutrophil transcriptome, which lead to abnormal neutrophil aging (21). Furthermore, we found that GCs trigger Rho GTPase signaling pathways. Phosphoproteomics also revealed the activations of ROCK and PAK protein kinases, which were associated with significant increases in actin cytoskeletal proteins and enrichment of Reactome pathway of Rho GTPase signaling at the proteome level. Lastly, we provide evidence that RIPOR2 is a novel target protein of DEX and PRED treatments. RIPOR2 protein levels were markedly elevated after GC treatments, accompanied by an even greater increase in relative phosphorylation levels at multiple serine sites (21, 489, 573, 585, and 588). Interestingly, RIPOR2 protein level was synergistically increased by the combination of DEX and PAKi treatment using PF-3758309. This combination treatment also enhanced GR phosphorylation and PINK1 protein level in immunoblotting analysis. Together, we speculate that GC-mediated activation of PAK is associated with RIPOR2 regulation that results in Rho GTPase signaling pathways.

Neutrophils are more transcriptionally inactive than other hematopoetic cells with lower total RNA content (58). Short lifespan of neutrophils remains a major obstacle for interrogated research, together with its minor contribution to adaptive immunity. Therefore, the anti-inflammatory effects of GCs have been less studied in neutrophils compared to other immune cell types, such as LPS-induced macrophages (59, 60), T cells (61), and natural killer cells (62). Nevertheless, investigating the plasticity and heterogeneity of neutrophils remains essential under disease and homeostatic status, given their critical roles in immunopathophysiology (63). The complexity of biological machinery increases by post-transcriptional processes including proteolytic cleavage, protein turnover, and variation in translation rates (64). In this context, MS-based proteomics offers a powerful approach for comprehensive analysis of neutrophils by utilizing quantification of global protein expressions and PTMs.

To overcome the limitations of short lifespan of neutrophils, we implemented an *in vitro* model by differentiation of a human promyelocyte cell line HL-60 using DMSO in medium containing serum, which has been reported to have the best differentiation potential for reactive oxygen species (ROS) formation and NET release (65). We confirmed neutrophil-like characteristics in the differentiated NLCs by cell cycle arrest and detection of mature neutrophil-specific cell surface markers. Yet, the principal component analysis of NLCs showed higher similarities with parental HL-60 cells than neutrophils (Fig.S6E). Another drawback of the *in vitro* differentiation system is the limited capacity to examine key neutrophil functions, likely due to the incomplete priming of NLCs to a fully functional and physiologically active state. We were unable to examine the release of NETs despite a robust upregulation of neutrophil elastase (*ELANE*) expression following GC treatments. Additionally, while GCs regulated the expression of actin-associated proteins, no significant changes were observed in cell migration across transwell membranes (data not shown).

GC treatments trigger TLR and MAPK cascades in NLCs and we collectively suggest that GCs stimulate abnormal neutrophil activation and aging. Neutrophil activation and aging share pathways associated with NOD-like receptors, NF-κB, and TLRs as they receive inflammatory signals for activation while they age in circulation. However, it is crucial to understand that the two processes are independent in terms of gene set enrichment within MAPK signaling pathways and cytokine/chemokine secretion (8). The *in vitro* setting lacks complex pathogenic signals that happen in the blood stream and does not allow separation of activation from aging. Nevertheless, the *in vitro* differentiation system allows unrestricted access to materials for deep proteome and phosphoproteome profiling that enables comprehensive molecular signaling pathway analysis. Using this approach, we provide proteome profiles of human neutrophils from buffy coats, HL-60 cells, and NLCs, which could be valuable resources to further understand the differentiation system. Moreover, we used two different synthetic GCs used in the clinics, DEX and PRED, to gain deeper insights into GR modulation. We focused our analysis on DEX because it is five to six times more potent with longer half-life than PRED (66), which was confirmed by more significant regulation of GR in proteome and phosphoproteome profiles.

We found multiple novel targets of GCs that have not been reported before. RIPOR2, first cloned in DMSO-differentiated HL-60 cells (67), facilitates directional migration of neutrophils by binding to RhoA at the leading edge. The phosphorylation of RIPOR2 results in relocalization of RIPOR2 from the plasma membrane to the cytosol, thereby disrupting the interaction of RIPOR2 with RhoA. This is mediated by the interaction with 14-3-3, which is a key modification for Rho-inhibitory function (57, 68). We provide a new insight behind RIPOR2-mediated pathway by showing that increased GR phosphorylation can promote RIPOR2 expression and its phosphorylation. Ultimately, GR may be an important factor that drives the front-signal-dependent polarization in neutrophils. FLT3 is another interesting target of GCs. FLT3 is a receptor tyrosine kinase that is often mutated in acute myeloid leukemia patients. Activating FLT3 mutations leads to overexpression of GR and increases sensitivity for the combination treatment of FLT3 inhibitor and GCs (69, 70). Our IMAC-based method for phosphopeptide-enrichment is abundance biased focused on serine (Fig.S6F) and regulation of FLT3 tyrosine phosphorylation target substrates was not observed. Still, based on previously reported links between FLT3 and GCs as well as our data, it is worth investigating further. Lastly, it is important to note that GCs drive proteasomal degradation of GR through its hyperphosphorylation (33). This results in significant time-dependent downregulation of GR expression in proteome. Further investigation is required to examine how GR turnover affects global proteome and phosphoproteome of GC-treated neutrophils.

In summary, this study provides valuable insights into GC-mediated signaling pathways of neutrophils driven by protein turnover and modifications which could be used to better understand the function of GCs in the human innate immune system. We also demonstrate that GC receptor could be an interesting target to study neutrophil polarization, activation, and aging.

## Conclusions

Neutrophil research has been neglected despite its critical role in the human immune system. In this study, we performed quantitative MS experiments in NLCs and showed unconventional functions of GCs. We first verified that NLCs share features with neutrophils isolated from human blood by quantifying their proteome profiles. We then conducted an extensive time-course treatment of GCs in NLCs, highlighting that GCs drive cell activation and aging. GC-regulated proteins and phosphosites were significantly enriched in TLR and Rho GTPase pathways. Proteome and phosphoproteome profiles of GC-treated neutrophils provided new insights into how GCs can affect the innate immune system.

## Methods

### Cell culture, differentiation, and treatment

HL-60 cells were cultured at 37°C with 5% CO_2_ in RPMI 1640 (11875093; Gibco^TM^) supplemented with 10% fetal bovine serum (FBS) and penicillin/streptomycin. Cells were tested for mycoplasma regularly. HL-60 cells were differentiated into NLCs by incubation using 1.25% DMSO (D2650; Sigma-Aldrich) for four days (65, 71). Dexamethasone (D4902; Sigma-Aldrich), prednisolone (P6004; Sigma-Aldrich), and mifepristone (475838; Sigma-Aldrich) were reconstituted in DMSO and added to cell culture medium at 1 µM concentration, unless otherwise indicated.

### Neutrophil isolation from buffy coats

Buffy coats were kindly provided by the National Center for Cancer Immune Therapy (CCIT-DK) at Herlev Hospital. Neutrophils were isolated using a previously published method (72). In short, buffy coats were separated by LeucoSep^TM^ polypropylene tubes (10349081; Greiner) and red blood cells (RBCs) were lysed with 10x RBS lysis buffer (420302; BioLegend). Isolated neutrophils were either lysed fresh or treated with 0.1% DMSO or GCs at 37°C, 5% CO_2_ in RPMI 1640/10% FBS.

### Human protein atlas

Normalized RNA levels (nTPM) for GR (*NR3C1*) and RIPOR2 were obtained from HPA tissue dataset and HPA immune cell dataset of the Human Protein Atlas (https://www.proteinatlas.org/).

### Immunoblotting

After 1.25% DMSO incubation, NLCs were washed 2x with PBS and seeded in 6-well plates (2 x 10^6^ cells / well), followed by indicated concentration of GCs for indicated time. After treatment, cells were washed 2x with PBS and extracted with Laemmli buffer (2% SDS, 150 mM NaCl, 50 mM Tris pH 8.5), then lysed by sonication at 20% amplitude. Protein concentration was measured using Pierce^TM^ BCA protein assay kits (10741395; Thermo Fisher Scientific). 20-30 ug proteins were prepared with 4x LSD sample buffer (11549166; Invitrogen^TM^) and loaded to 4-12% Bis-Tris gels. 10 µl of SeeBlue^TM^ plus2 pre-stained protein standard (LC5925; Invitrogen^TM^) was loaded with samples. Proteins were separated by SDS-PAGE and transferred to nitrocellulose membranes. Membranes were blocked with 5% skim milk or 3% BSA, incubated with primary antibodies overnight at 4°C, and incubated with HRP-conjugated rabbit/mouse secondary antibodies for 1 hour at room temperature. Membranes were developed using ECL chemiluminescent substrate reagent kit (10348463; Invitrogen^TM^). The following primary antibodies were purchased from Cell Signaling Technology: p-GR (Ser211)(#4161), p-ERK1/2 (Thr202/Tyr204)(# 9106), t-ERK1/2 (#4695), p-p38MAPK (Thr180/Tyr182)(# 9211), t-p38MAPK (#9212), p-AKT (Ser473)(# 9271), t-AKT(#9272), and PINK1 (#6946). GR (sc393232), Vinculin (sc-25336 HRP), α-tubulin (sc-32293 HRP), and GAPDH (sc-365062 HRP) were purchased from Santa Cruz Biotechnology. Other primary antibodies used were Rac2 (ab211162; abcam), Ras (05-516; Millipore), and RIPOR2 (17015-1-AP; Proteintech).

### Cell viability assay

Cell viability was measured using CCK-8 assay. Cells were seeded in 96-well plates (1.5 x 10^4^ cells for HL-60 cells and 6 x 10^4^ cells for NLCs) containing 130 µL RPMI 1640/10% FBS medium. After treatment with DEX for indicated incubation time, 13 µL of WST-8 (CK04-13; Dojindo) was added to each well. After 3 hours, absorbance was measured at 450 nm using multi-mode microplate reader (FLUOstar Omega; BMG Labtech).

### Flow cytometry

Cells were washed 2x with ice-cold PBS/10% FBS. For apoptosis analysis, cells were resuspended in 100 µL of 1x binding buffer, then stained with 5 µL of propidium iodide (PI) and 0.5 µL of annexin V (556547). For cell cycle analysis, cells were stained with Fixable Viability Stain (FVS) 780 (565388), fixed with 70% EtOH, and resuspended with PI/RNase staining buffer (550825). For surface marker staining, we followed OMIP-100 method (29). Cells were stained with FVS780, blocked with Human TruStain FcX^TM^ (422302; BioLegend), and incubated with desired conjugated antibodies for 20 min: CD15 (323044), CD45 (566961), CD184 (562448). All stains were purchased from BD Biosciences. Samples were measured with Cytek Aurora (Cytek Biosciences) and analyzed with FCS Express 7 (De Novo Software).

### Cell lysis and protein digestion for mass spectrometry

#### Neutrophils isolated from buffy coats

Neutrophil samples were prepared using a previously published method (73). Cells were lysed with boiling 5% SDS buffer (5 mM TCEP, 10 mM CAA, 100 mM Tris pH 8.5) and incubated at 95°C for 10 min shaking at 1400 rpm. Benzonase was added and incubated at 37°C for 15 min. 3 µg of proteins were digested using PAC method (74) with MagReSyn Hydroxyl beads (Resyn Biosciences) and an enzyme:protein (µg) ratio of 1:200 for trypsin (T6567; Sigma-Aldrich) and 1:200 for lys-C (129-02541; FUJIFILM Wako Pure Chemical Corporation) in 50 mM triethyl ammonium bicarbonate. Digested peptides were directly loaded to Evotips using Opentrons OT-2 PAC digestion protocol available at https://www.evosep.com/support/automation-opentrons-ot2/.

#### NLC lysis with SDS buffer

DEX and PRED time-course treated proteome and phosphoproteome were prepared with SDS lysis buffer. Cells were washed 2x with ice-cold PBS containing phosphatase inhibitors (2 mM sodium orthovanadate, 1 mM β-glycerophosphate, 1 mM NaF). Cells were lysed with boiling 5% SDS buffer (5 mM TCEP, 10 mM CAA, 100 mM Tris pH 8.5), incubated at 95°C for 10 min shaking at 1400 rpm, and sonicated at 40% amplitude for 5 sec x2. 500 µg of proteins were PAC digested using KingFisher (Thermo Fisher Scientific) and eluted in 50 mM ammonium bicarbonate containing 2 µg trypsin and 1 µg lys-C. After overnight digestion, samples were desalted by Sep-Pak C18 96-well plate (186003966; Waters) and eluted with 80% acetonitrile (ACN). 1% of samples were loaded onto Evotips for single-shot proteome analysis, and the rest were used for phosphopeptide enrichment.

#### NLC lysis with GdnHCl buffer

HL-60 cells and NLCs treated with DEX and MIF for 24 hours were prepared with guanidinium chloride (GdnHCl) lysis buffer. Cells were lysed with 6 M GdnHCl buffer (5 mM TCEP, 10 mM CAA, 50 mM Tris pH 8.5) and sonicated at 30% amplitude for 3 sec x2. Protein concentration was measured by Bradford assay. 100 µg of proteins were diluted to 1.5 M GdnHCl and incubated overnight at room temperature with 0.5 µg lys-C in 50 mM Tris pH 8.5. On the next day, samples were acidified with 10% TFA and desalted with 8-layers of C18 StageTips as previously described (75). Samples were eluted with 30% ACN in 0.1% formic acid, speedvac dried, and reconstituted with 50 mM ammonium bicarbonate containing 0.5 µg trypsin. After overnight incubation at room temperature, samples were speedvac dried and reconstituted in 20 µL of 0.1% formic acid. 10% of samples were run on LC-MS/MS.

#### Phosphopeptide enrichment

After eluting SDS-lysed samples with 80% ACN following desalting by Sep-Pak, eluates were adjusted to 80% ACN/5% TFA/0.1M glycolic acid (GA). Samples were incubated with 12 µL of Ti-IMAC beads (MagResyn) and washed with 80% ACN/5%TFA/0.1M GA, 80% ACN/1% TFA, and 10% ACN/0.2% TFA using KingFisher. Phosphoenriched peptides were eluted from beads with 1% NH_3_OH, acidified with 10% TFA, and loaded onto Evotips for LC-MS/MS analysis.

### Mass spectrometry analysis

#### Evosep One LC – Orbitrap Astral MS

SDS-lysed samples were analyzed on Evosep One LC system coupled with Orbitrap Astral MS (Thermo Fisher Scientific) in data independent acquisition (DIA) mode. Samples were separated with 15 cm x 150 µM performance column (EV1137; Evosep) at a gradient of 30 samples per day. Full scan orbitrap resolution was set to 240,000 in positive ion mode, ion transfer temperature (ITT) 275°C, RF lens 40%, normalized AGC target 500%, and isolation window (m/z) of 2. For single-shot proteome, full scan range (m/z) was set to 380-980 with maximum injection time of 3 ms. For phosphoproteome, full scan range (m/z) was set to 480-1080 with maximum injection time of 30 ms.

#### Vanquish Neo LC – Orbitrap Astral MS

GdnHCl-lysed samples were analyzed on Vanquish Neo UHPLC system (Thermo Fisher Scientific) coupled with Orbitrap Astral MS in DIA mode. Samples were separated with home-pulled 15 cm x 75 µM column packed with ReproSil-Pur 120 C18-AQ 1.9 mm beads (Dr. Maisch) by a gradient of 80% ACN ranging from 5%-90% for 17 min. Full scan orbitrap resolution was set to 240,000 in positive ion mode, ITT 275°C, RF lens 50%, normalized AGC target 250%, and isolation window (m/z) of 4. Full scan range (m/z) was set to 300-1000 with maximum injection time of 50 ms.

#### Raw file analysis

All MS raw files were analyzed with Spectronaut (Biognosys) v19 using human UniprotKB fasta protein sequence database (reviewed in May 2024) and common contaminants in a library-free mode (direct DIA+). For fixed modification, cysteine carbamidomethylation was selected. For variable modification, protein N terminus acetylation and methionine oxidation were selected. Quantification was calculated with cross-run normalization disabled. Phosphoproteome was searched with an additional variable modification of serine, threonine, and tyrosine phosphorylation. Minimum localization probability was set to 0.75.

### Bioinformatics

Using Perseus v2.0.11, protein groups and phosphosites were kept if at least four valid values were present per condition. Label-free quantifications were log_2_ transformed, then imputed with width 0.15 and down shift 2 in total matrix mode. Data was further analyzed using R (v4.4.1) and RStudio (v2024.04.2). MS quantifications were median normalized and statistical analysis was performed using one-way analysis of variance (ANOVA) followed by Tukey’s Honestly Significant Difference (HSD) post hoc test using R or one-way ANOVA with Dunnette’s multiple comparisons test using GraphPad Prism 10. Differential expressions were considered statistically significant at adjusted *p*-value < 0.05. Soft clustering of significant protein groups was performed using Mfuzz clustering package with fuzzification factor *m* = 1.5 and cluster number 8. Volcano and scatter plots were created using ggplot2 package. Kinase-substrate datasets were obtained from PhosphoSitePlus and SIGNOR 3.0. Protein networks and pathway enrichments were analyzed using STRING (https://string-db.org/) and Cytoscape (v3.10.0) with whole genome as background. Kinase activity was analyzed using RoKAI App (https://rokai.io/) with minimum substrate of 5 and absolute z score of 2.

## Figure Legends

**Supplementary Figure 1.**
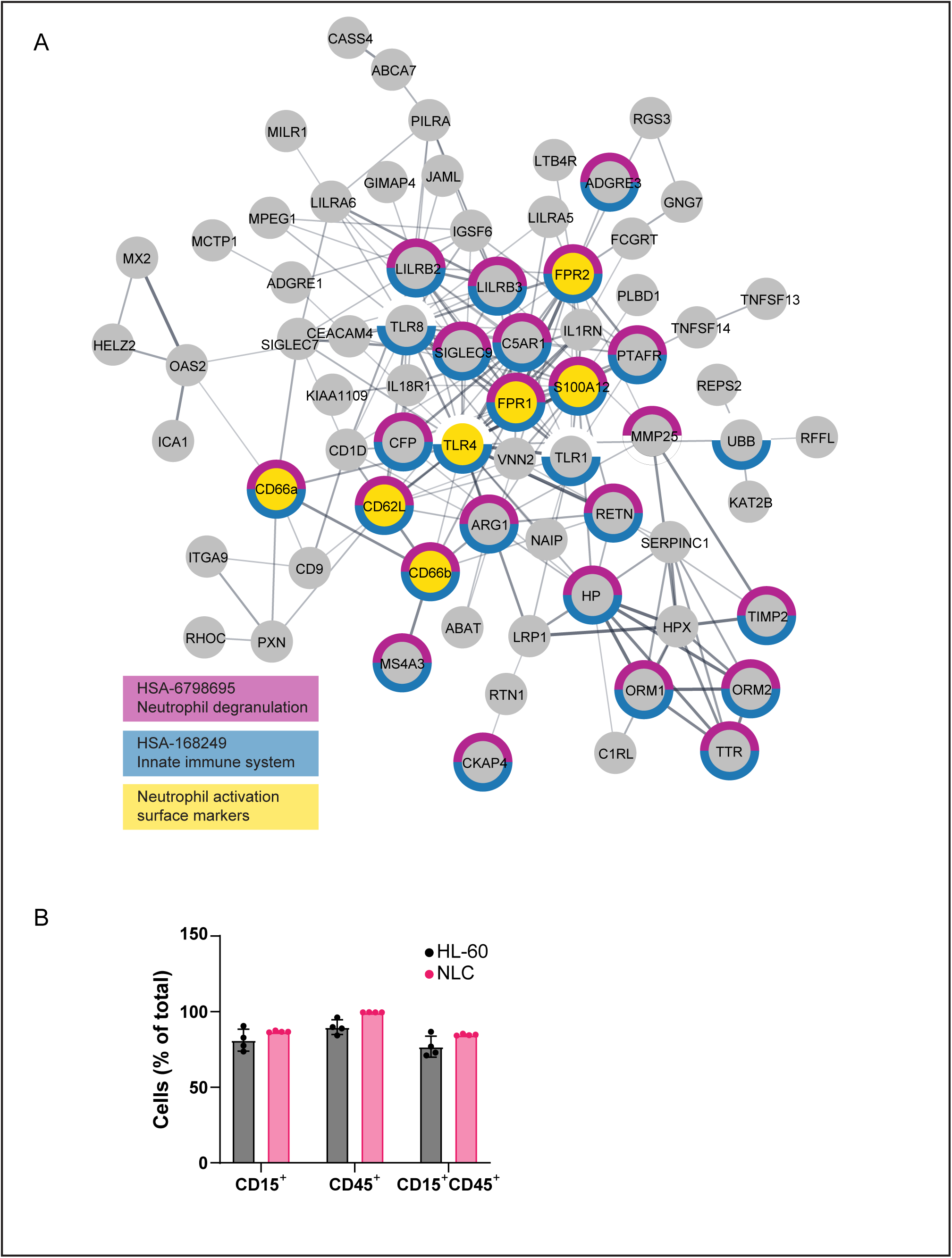
A) String network of 144 proteins uniquely quantified in NLCs and neutrophils. Proteins with specific functions are highlighted: neutrophil degranulation (magenta), innate immune system (blue), and neutrophil activation surface markers (yellow). B) Flow cytometry analysis of CD15 and CD45 staining in HL-60 cells and NLCs. Bar plot represents CD15^+^ and CD45^+^ cell population from gated live cells.

**Supplementary Figure 2.**
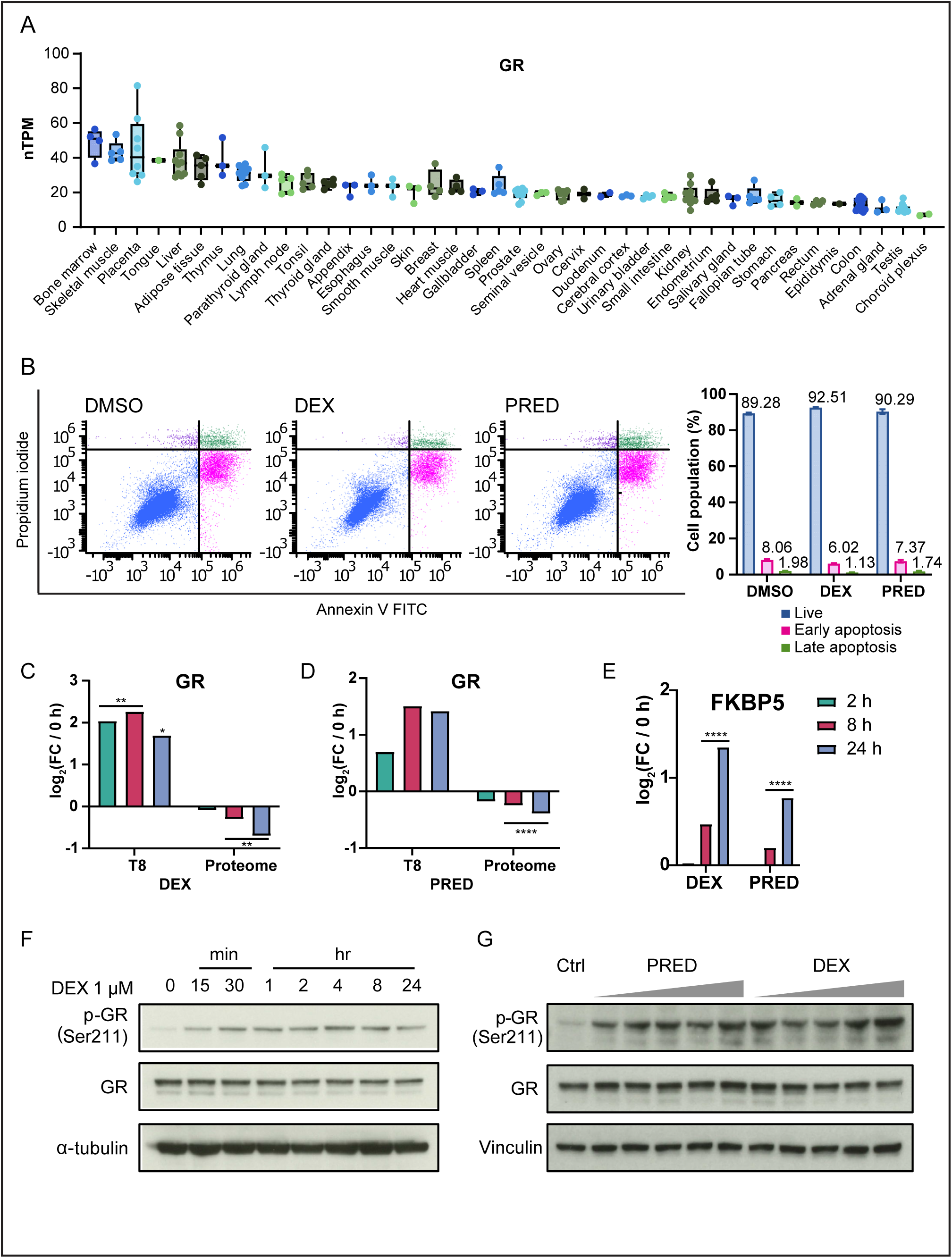
A) GR mRNA levels represented as normalized transcript per million (nTPM) in human tissues. RNA-seq data was acquired from the Human Protein Atlas (HPA). B) Apoptosis assay of NLCs after 24 hours treatment of DEX and PRED. Cells were stained with propidium iodide and annexin V, then analyzed with flow cytometry. Bar plot represents live, early apoptosis, and late apoptosis cell populations from gated live cells. C-D) Bar plots showing log_2_ transformed fold changes of GR phosphorylation at Thr-8 and total form after DEX (C) and PRED (D) treatments. E) Bar plots showing log_2_ transformed fold changes FKBP5 protein expression after DEX and PRED treatments. One-way ANOVA with Tukey’s HSD correction was used. F-G) Immunoblotting analysis of GR Ser-211 phosphorylation and total form after time-dependent treatment of 1 µM DEX (F) and dose-dependent treatment of DEX and PRED for 24 hours (0.1, 0.5, 1, 5, 10 µM) (G).

**Supplementary Figure 3.**
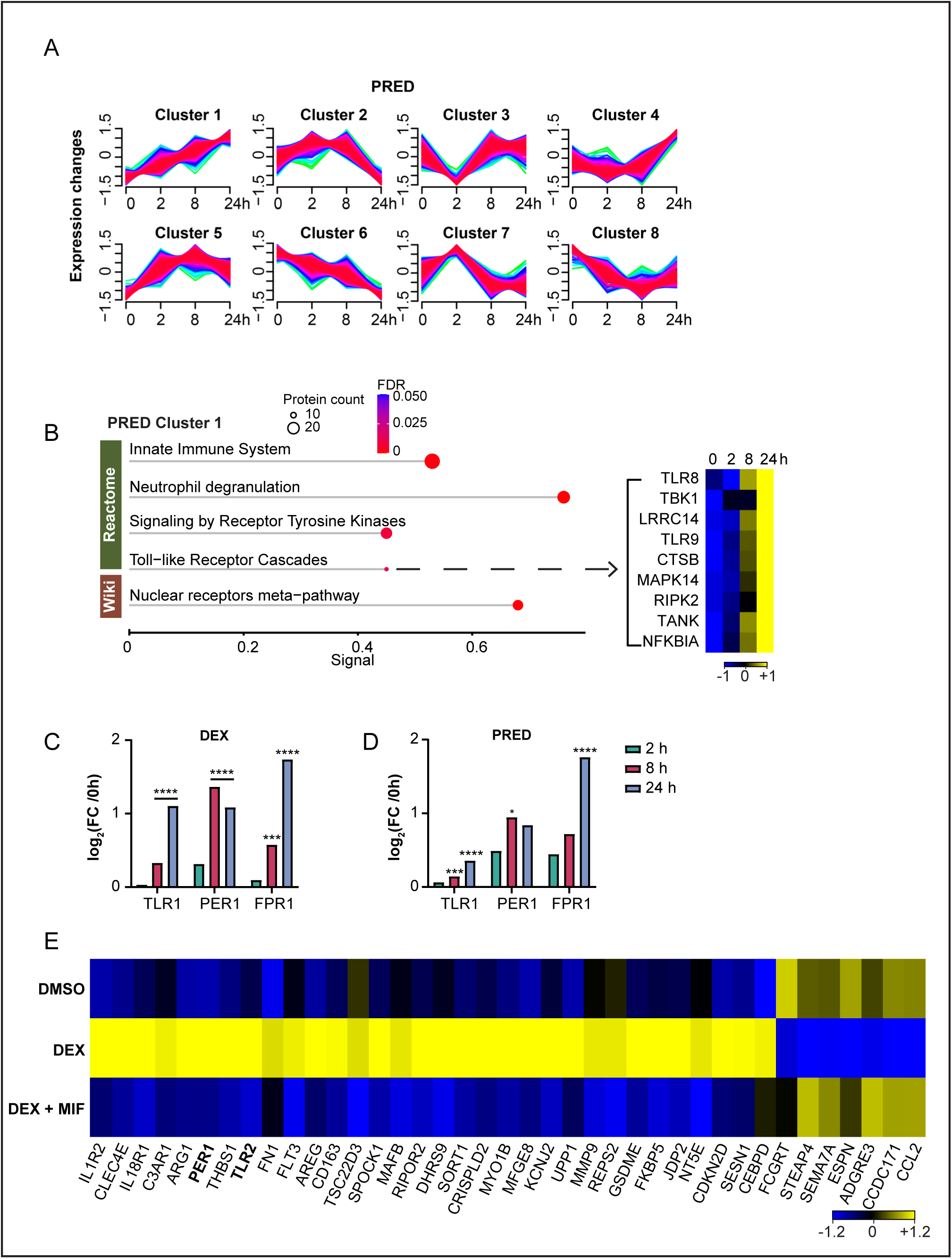
A) Soft clustering analysis with statistically significant proteins (adjusted *p*-value < 0.05, one-way ANOVA with Tukey’s HSD correction) after PRED treatment in NLCs. Mfuzz clustering package was used with fuzzification factor *m* = 1.5 and cluster number 8. B) Functional enrichment of proteins in cluster 1 using String analysis after PRED treatment. Dot size represents the number of proteins and dot color represents FDR values. Heatmap represents proteins in TLR cascades with z-scored log_2_ transformed label-free quantification, colors showing z scores between −1 and 1. C-D) Bar plots showing log_2_ transformed fold changes of TLR2, PER1, and FPR1 protein expressions after DEX (C) and PRED (D) treatments. E) Heatmap representing proteins with statistically significant differences (adjusted *p*-value < 0.05 and log_2_FC > |1|) in either DEX or DEX+MIF treatment compared to DMSO control. One-way ANOVA with Tukey’s HSD correction was used. Log_2_ transformed label-free quantification was z-scored and represented in colors ranging from −1.2 to 1.2

**Supplementary Figure 4.**
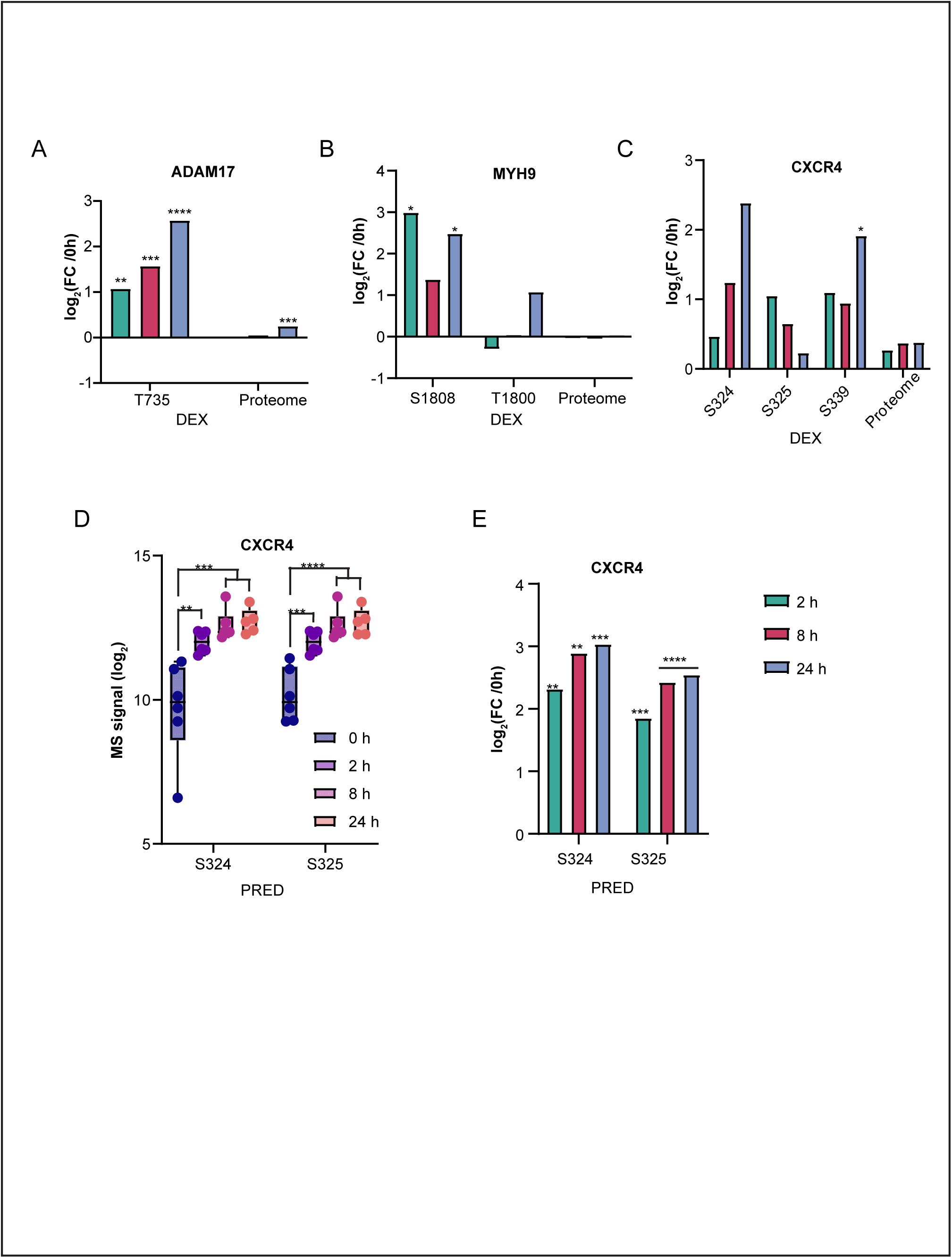
A-C) Bar plots showing log_2_ transformed fold changes of ADAM17 (A), MYH9 (B), and CXCR4 (C) phosphorylation and protein expressions after DEX treatment. D) Box plots showing individual log_2_ transformed label-free quantification of CXCR4 serine phophsites after PRED treatment. E) Bar plots showing log_2_ transformed fold changes of CXCR4 serine phosphorylation after PRED treatment. One-way ANOVA with Dunnette’s multiple comparisons test was used for box plot analysis and one-way ANOVA with Tukey’s HSD correction was used for bar plot analysis.

**Supplementary Figure 5.**
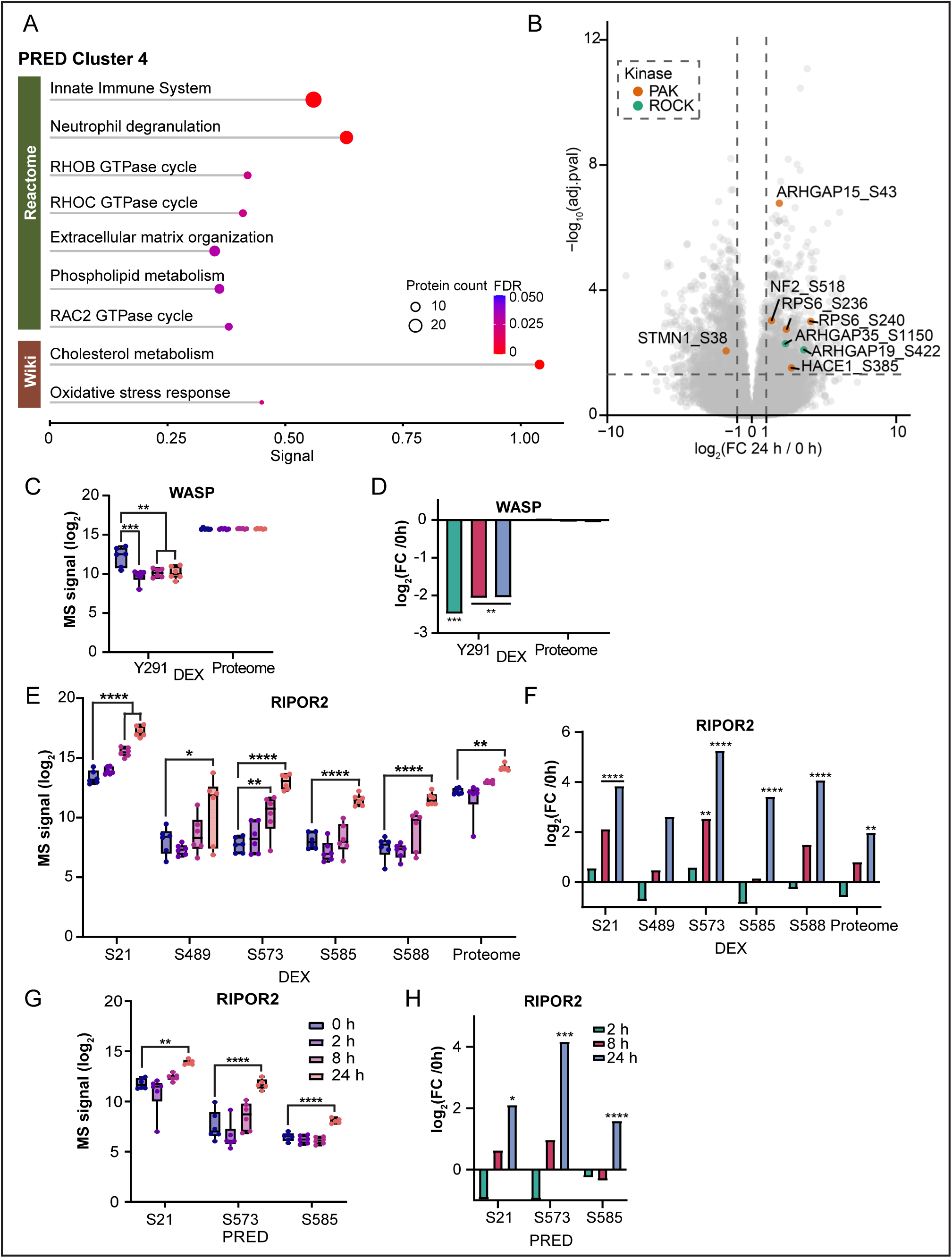
A) Functional enrichment of proteins in cluster 4 using String analysis after PRED treatment. Dot size represents the number of proteins and dot color represents FDR values. B) Volcano plot comparing phosphorylation expressions of DEX 24 hours treatment versus 0 hour control in NLCs using one-way ANOVA with Tukey’s HSD correction. Dots highlighted in orange are substrates of PAKs and dots in green are substrates of ROCKs. C, E) Box plots showing individual log2 transformed label-free quantification of WASP (C), RIPOR2 (E) phosphorylation and protein expressions after DEX treatment. D, F) Bar plots showing log_2_ transformed fold changes of WASP (D), RIPOR2 (F) phosphorylation and protein expressions after DEX treatment. G) Box plots showing individual log_2_ transformed label-free quantification of RIPOR2 serine phophsites after PRED treatment. H) Bar plots showing log_2_ transformed fold changes of RIPOR2 serine phosphorylation after PRED treatment. One-way ANOVA with Dunnette’s multiple comparisons test was used for box plot analysis and one-way ANOVA with Tukey’s HSD correction was used for bar plot analysis.

**Supplementary Figure 6.**
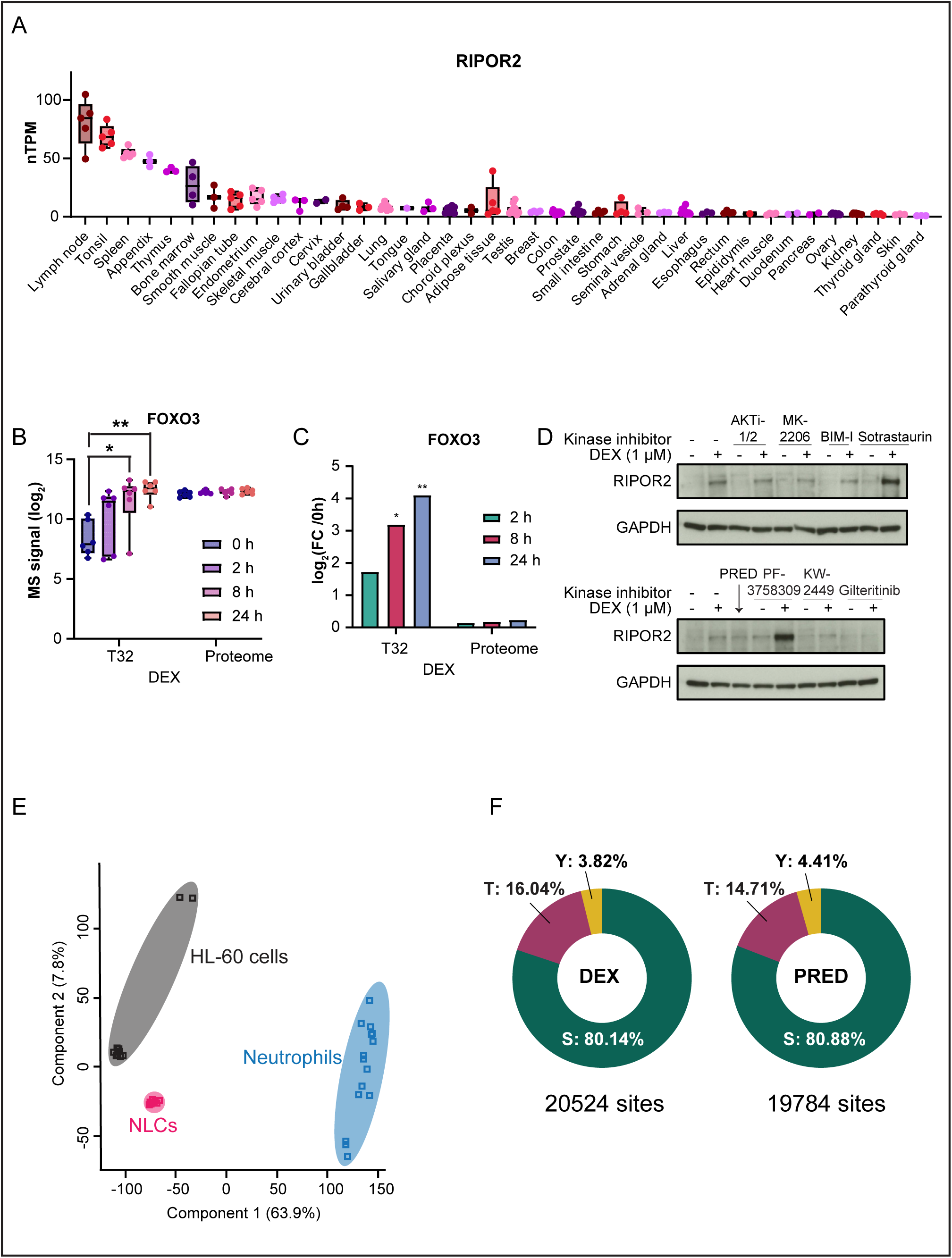
A) RIPOR2 mRNA levels represented as normalized transcript per million (nTPM) in human tissues. RNA-seq data was acquired from the Human Protein Atlas (HPA). B) Box plots showing individual log_2_ transformed label-free quantification of FOXO3 phosphorylation and protein expressions after DEX treatment. C) Bar plots showing log_2_ transformed fold changes of FOXO3 phosphorylation and protein expressions after DEX treatment. One-way ANOVA with Dunnette’s multiple comparisons test was used for box plot analysis and one-way ANOVA with Tukey’s HSD correction was used for bar plot analysis. D) Immunoblotting analysis of kinase inhibitors in combination with DEX treatment for 24 hours in NLCs. E) Principal component analysis (PCA) of neutrophils, HL-60 cells, and NLCs. Component 1 (x-axis) and component 2 (y-axis) were used. F) Pie charts showing the proportion of serine, threonine, and tyrosine in phosphoproteome after DEX and PRED treatments in HL-60 cells and NLCs.

